# Set1 regulates telomere function via H3K4 methylation-dependent and independent pathways and calibrates the abundance of telomere maintenance factors

**DOI:** 10.1101/2022.06.08.495211

**Authors:** Meagan Jezek, Winny Sun, Maraki Y. Negesse, Zachary M. Smith, Alexander Orosz, Erin M. Green

## Abstract

Set1 is an H3K4 methyltransferase which comprises the catalytic subunit of the COMPASS complex and has been implicated in transcription, DNA repair, cell cycle control, and numerous other genomic functions. Set1 also promotes proper telomere maintenance, as cells lacking Set1 have short telomeres and disrupted subtelomeric gene repression; however, the precise role for Set1 in these processes has not been fully defined. In this study, we have tested mutants of Set1 and the COMPASS complex which differentially alter H3K4 methylation status and attempt to separate catalytic and non-catalytic functions of Set1. Our data reveal that Set1-dependent subtelomeric gene repression relies on its catalytic activity towards H3K4, whereas telomere length is regulated by Set1 catalytic activity but likely independent of the H3K4 substrate. Furthermore, we uncover a role for Set1 in calibrating the abundance of critical telomere maintenance proteins, including components of the telomerase holoenzyme and members of the telomere capping CST (Cdc13-Stn1-Ten1) complex, through both transcriptional and post-transcriptional pathways. Altogether, our data provide new insights into the H3K4 methylation-dependent and independent roles for Set1 in telomere maintenance in yeast and shed light on possible roles for Set1-related methyltransferases in other systems.

## Introduction

As the catalytic component of the COMPASS complex, the methyltransferase Set1 deposits mono-, di-, and tri-methylation onto histone H3 lysine 4 (1–4). In yeast, H3K4me3, me2, and me1 are found at gene promoters, gene bodies, and 3’ regions of actively transcribed genes, respectively (5–7). Interestingly, in the absence of Set1, genome-wide gene expression analyses have shown that the majority of differentially expressed genes are up-regulated (8–13), indicating that a primary outcome of depleted Set1 and H3K4 methylation is gene derepression. The genes that rely on Set1 for repression are predominantly lowly expressed and are found in genomic regions with low transcriptional activity, such as Ty elements, rDNA, meiotic differentiation genes, and telomeres (13–18).

One of the earliest described phenotypes in yeast cells lacking Set1 was the loss of subtelomeric gene silencing (2,14,19). *set1Δ* cells also have short telomeres and disrupted telomere clustering at the nuclear periphery (2,14,20,21). Given the low levels of H3K4 methylation at telomeric regions, it was proposed that Set1 indirectly regulates subtelomeric gene silencing, and likely telomere length, through euchromatic H3K4me3, which is thought to prevent redistribution of the silencing protein Sir2, the histone deacetylase (HDAC) of the SIR complex, from subtelomeres to internal chromosomal sites (10,22). However, subsequent chromatin immunoprecipitation and genetic studies indicate that subtelomeric gene repression by Set1 predominantly occurs through a Sir-independent mechanism (9,12,23,24). These observations are consistent with recent understanding of SIR-mediated silencing at subtelomeres, which revealed largely noncontiguous subtelomeric regions reliant on the SIR complex for silencing, whereas Set1 appears to promote silencing in larger, contiguous regions and at a higher number of subtelomeres (12,25). These observations indicate that the precise role for Set1 in maintaining functional telomere-linked properties such as gene repression, telomere length, and nuclear position remains obscure. Interestingly, in previous work, our lab has shown that the *set1Δ* transcriptome is highly similar to mutants lacking telomere maintenance factors, particularly the telomerase subunit, Est3 (12). This further suggests that Set1 may have a more direct role in telomere regulation than previously appreciated.

Although most known functions of Set1 have been attributed to methylation of H3K4, there is growing evidence for H3K4me-independent functions for Set1. For example, two other substrates have been described for Set1: the kinetochore protein, Dam1 (26), and another histone modification at H3K37, for which Set1 promotes the methylation with Set2 (27). Although not extensively documented for yeast Set1, some orthologs of Set1 have been demonstrated to perform non-catalytic functions as well (28–31). The discovery of these additional substrates and functions implicates Set1 in genome regulation roles separate from its H3K4 methyltransferase activity.

In wildtype cells, multiple protein complexes bind telomere ends and coordinate telomere capping and elongation. These include the Cdc13-Stn1-Ten1 (CST) capping complex and telomerase, composed of the catalytic subunit Est2, regulatory subunits Est1 and Est3, and the RNA component *TLC1* (32–34). These and other proteins are critical to telomere health and numerous mechanisms appear to control their precise abundance in cells, including at the level of transcription, translation, and regulated degradation (35–46). These studies have revealed that changes in their stoichiometry alter telomere length and stability, indicating that a precise and well-defined balance in telomerase and CST components is critical to telomere maintenance.

In this study, we assayed mutations in Set1, the COMPASS complex, and at H3K4 to assess the specific contribution of H3K4 methylation by Set1 to subtelomeric gene repression and telomere length maintenance. Our data indicate that the catalytic core of Set1 is required both for its role in gene repression and telomere length; however there are likely H3K4 methyl-independent mechanisms required for the regulation of telomere length. Furthermore, our data identify a role for Set1 and H3K4 methylation in modulating the abundance of telomere maintenance proteins, including components of telomerase and the CST complex, through both transcriptional and post-transcriptional pathways. Altogether, these data provide new insights into how Set1 contributes to telomere maintenance and is likely applicable to related proteins in other organisms.

## Materials and methods

### Yeast strains and growth conditions

Yeast strains, plasmids, and primers used in this study are listed in **Table S1, S2,** and **S3**, respectively. Gene knockouts, incorporation of epitope tags, and plasmid construction and transformation were performed using standard methods (13,47–50). The *FLAG-SET1* allele was constructed by targeting of a 2xFLAG-containing PCR cassette upstream of the *SET1* coding sequence, as described (51). This sequence was PCR-amplified from genomic DNA, along with the *SET1* promoter and 3’UTR, and cloned into pRS316 for generating mutants in a plasmid-based expression system. Amino acid substitutions and domain deletions were made using either Gibson assembly or the Q5 Site-Directed Mutagenesis Kit (NEB). Isogenic single and double mutant strains were generated via haploid mating, sporulation, and tetrad dissection. Genotypes were confirmed using PCR (endogenous mutations) or plasmid sequencing. Standard liquid and solid growth media were used (YPD and SC-dropout). The *CDC13* temperature-sensitive mutant strain was a gift from David Lydall (Newcastle University) (35).

### RNA extraction and RT-qPCR to measure mRNA abundance

RNA was extracted from 1.5 mL of logarithmically growing cells (OD_600_ ~0.6-0.8) using the Masterpure Yeast RNA Purification kit (Epicentre). Residual genomic DNA was removed using the Turbo DNA-free kit (Ambion). cDNA was synthesized using qMax cDNA Synthesis Kit (Accuris) with 1 μg template RNA. Quantitative PCR (qPCR) was performed to assess mRNA transcript levels with qMax Green Low Rox qPCR Mix (Accuris), 0.5 μL of cDNA and 20 μM of forward and reverse gene specific primers. Amplification was performed using a CFX384 Real-time Detection System (Bio-Rad). Each reaction was performed in technical triplicate, and a minimum of three biological replicates. mRNA abundance was determined relative to the control gene *TFC1* and displayed as fold change (or relative expression) compared to wildtype. To measure mRNA degradation, cells were treated with 10 μg/mL thiolutin (ENZO Life Sciences) and harvested at 10 and 20 minutes after treatment. mRNA abundance was determined relative to the control gene *sCR1* and normalized to the zero time point. All qPCR primers used are listed in **Table S3**.

### DNA preparation and Southern blots of telomere length

Telomere length was assessed using a non-radioactive Southern blotting protocol (52). Briefly, 10 mL of yeast culture was grown overnight at 30°C, harvested and pellets were resuspended in Yeast Breaking Buffer (10mM Tris-HCl pH 8.0, 100mM NaCl, 1mM EDTA pH 8.0, 1% SDS, 2% Triton X-100) and vortexed with acid-washed glass beads (Sigma-Aldrich) with phenol-chloroform-isoamyl alcohol (25:24:1) (VWR Scientific). After centrifugation, DNA was isolated from the supernatant via ethanol precipitation and treated with RNaseA. Genomic DNA was digested with XhoI (NEB) and subject to PCI extraction and ethanol precipitation. 40-60 μg of prepared DNA was loaded onto an 0.8% agarose gel (0.5% TBE) and run for 25hr at 100V. Ethidium bromide staining and imaging was used to observe DNA migration. Following denaturation and washing (52), DNA was transferred onto an N+hybond membrane (Amersham) prior to crosslinking. The blot was probed with 15 ng/ml biotinylated telomere probe (5’-biotin-CACACCCACACCCACACC-3’) at 65°C overnight. Nucleic acids on the blot were detected using a Chemiluminescent Nucleic Acid Detection Module Kit (ThermoFisher) per the manufacturer’s instructions and imaged using a LiCor C-Digit chemiluminescent imager.

### Yeast senescence assays

To obtain replicatively “young” cells for all appropriate genotypes, tetrad dissections of heterozygous diploid strains were used to generate isogenic haploid strains. Senescence assays were performed as described (53,54). Briefly, cells were grown on plates for three days and single colonies were used to inoculate 10 mL YPD which were grown overnight shaking at 30°C. Cell density was calculated using a hemocytometer and new cultures were diluted to 1×10^4^ cells/mL from the overnight cultures. After 24 hours, cell density was recorded and new cultures were inoculated at 1×10^4^ cells/mL. This procedure was repeated every 24 hours for up to 15 days. The number of population doublings per culture was calculated for each time point taken. Experiments were performed in biological triplicate.

### Yeast spot assays

Cells were grown at 30°C to saturation in 5 mL YPD and then diluted to 1.0 OD_600_ units. Ten-fold serial dilutions were plated on the appropriate media, including YPD, SC-URA, SC-URA + 100 mM hydroxyurea, or SC-URA + 10 mM caffeine. For temperature sensitive assays, plates were grown at the indicated temperatures and imaged up to 4 days. For DNA damage sensitivity assays, plates were grown at 30°C and imaged daily for up to 7 days.

### Immunoblotting

Lysates for immunoblotting were prepared using either 0.2M NaOH treatment or bead-beating in lysis buffer (50mM Tris-HCl pH 7.5, 150mM NaCl, 1mM EDTA, 1% NP-40) (12,49). Protein concentrations were determined by Bradford assay and normalized to load equivalent amounts on SDS-PAGE. Proteins were transferred to Immobilon-FL PVDF membrane (Fisher Scientific). Blots were probed overnight with primary antibodies, followed by incubation with the appropriate secondary antibody. Blots were imaged using an Odyssey CLx Scanner and processed using ImageStudio (LiCor). Total Protein Stain (LiCor) was used to determine total protein amounts for quantitation. Antibodies used in this study: c-Myc 9E10 (Novus Biologicals NB600-302SS), Monoclonal Anti-FLAG M2 (Sigma-Aldrich F1804), H3K4me3 (Active Motif 39159), H3K4me2 (Active Motif 39142), H3 (Abcam ab1791), H4 (Abcam ab31830), IRDye 680RD Goat anti-Mouse IgG (LiCor 926-68070), IRDye 800CW Goat anti-Rabbit IgG (LiCor 926-32211).

### Chromatin Immunoprecipitation

Chromatin immunoprecipitation was performed as previously described (12,48,55). All primers used for amplification of chromatin regions are listed in **Table S3**. The ratio of percent input for the histone mark (H3K4me3 or H3K4me2) is shown relative to percent input for histone H3.

## Results

### The Set1 SET domain maintains telomere length and subtelomeric gene repression

In the absence of Set1, cells display a number of telomere-related defects (2,12,14,19–21), including loss of subtelomeric gene repression as demonstrated by derepression of *TEL07L*-adjacent genes *COS12* and *YGL262W* (**Figure 1A**), and short telomeres (**Figure 1B**). In investigating other telomere-related phenotypes of *set1Δ* cells, we also observed a faster rate of senescence in cells lacking Set1 and the telomerase component *TLC1,* as compared to *tlc1Δ* single mutants (**Figure 1C**). These data show that the telomeres in *set1Δ tlc1Δ* cells are shortened critically to reach a crisis point, prompting cellular senescence. The *set1Δ tlc1Δ* cells are able to recover, however, and form survivors through a telomerase-independent mechanism, albeit at a slower rate than *tlc1Δ* cells (**Figure 1C**). We also found that loss of Set1 increases the temperature sensitivity of a strain carrying a temperature-sensitive mutation in *CDC13,* which encodes a member of the CST telomere capping complex (**Figure 1D**). While *cdc13-1* cells show almost no growth at 30°C and some growth at 27°C, the *set1Δ cdc13-1* mutants do not grow at 27°C, indicating that Set1 functions in a related pathway to Cdc13 in telomere function. This is consistent with our previous observations indicating a broader role for Set1 in telomere maintenance pathways (12), rather than a targeted function in controlling subtelomeric gene silencing.

**Figure 1.**
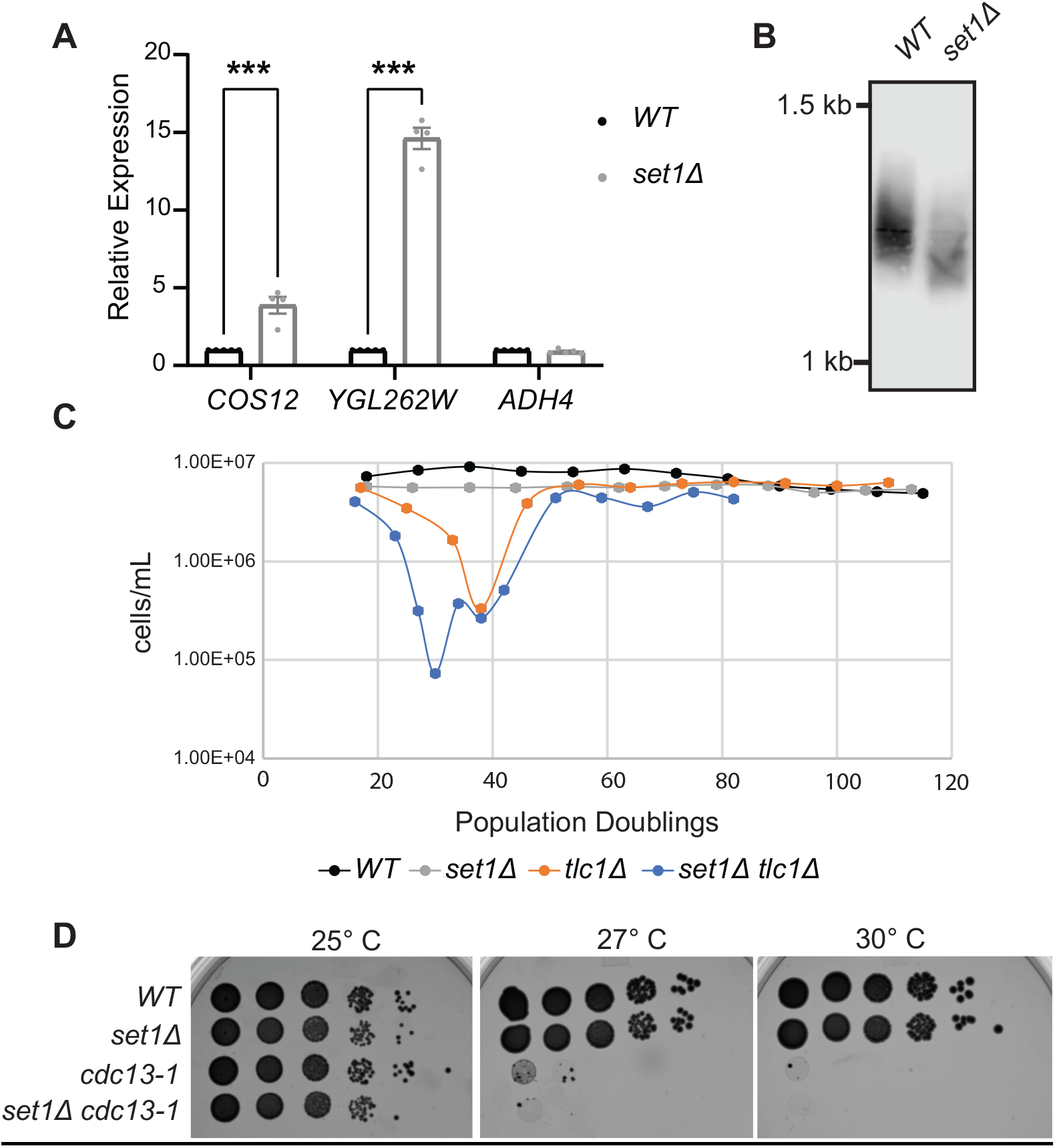
Loss of Set1 is linked to telomere defects. (**A**) RT-qPCR analysis of *TEL07L* genes *(COS12, YGL262W,* and *ADH4)* in wildtype *(WT)* and *set1Δ* cells. Expression normalized to the control gene *TFC1* and relative to *WT* is shown. Error bars represent standard error of the mean (SEM) for a minimum of three biological replicates. Significance was evaluated using two-way ANOVA and Turkey’s multiple comparisons test. *p*-values are indicated as follows: * < 0.05, ** < 0.01, *** <0.001. (**B**) Southern blot showing terminal telomere fragment molecular weight in *WT* and *set1Δ* cells. (**C**) Senescence rate of *WT*, *set1Δ, tlc1Δ,* and *set1Δ tlc1Δ* cells. At each time point, population doublings were calculated based on the cell density. The average of three biological replicates is shown and error bars indicate SEM. (**D**) Ten-fold serial dilutions of saturated cultures of the indicated strains were spotted on YPD and incubated at different temperatures prior to imaging.

To better define the molecular contribution of Set1 in preventing telomere-related defects, we monitored subtelomeric silencing and telomere length in mutants specifically lacking H3K4 methylation by carrying an allele expressing the *H3K4R* mutant as the sole copy of an H3-encoding gene. These cells showed similar derepression of subtelomeric genes as *set1Δ* cells (**Figure 2A**) and also a similar shortening in telomere length (**Figure 2B**). We combined the *H3K4R* mutant with *set1Δ* and also observed that the subtelomeric gene repression and telomere length were reduced to a similar extent in the double mutant compared to both single mutants (**Figures 2A** and **2B**). While this suggests an epistatic interaction between *SET1* and *H3K4*, as is expected based on their biochemical relationship, previous work has demonstrated that the Set1 protein is destabilized in the absence of H3K4 methylation in these mutants due to an autoregulatory feedback mechanism (56). Therefore it is possible that the phenotypes in these mutants are primarily indicative of a loss of Set1 alone. We attempted to further address whether the methyltransferase activity of Set1 is required for subtelomeric gene repression and telomere length maintenance by generating multiple catalytically inactive mutations in cells expressing *FLAG-SET1* from a plasmid under the control of the *SET1* promoter in *set1Δ* cells, including *H1017L, C1019A,* and *G951S* (14,57–59). While we did observe complete loss of H3K4 methylation in these mutants, we also found that their steady-state protein levels were much lower than wildtype FLAG-Set1 (**Figure 2D**), as previously-shown for some catalytically inactive mutants (56). As expected, the mutants all showed derepression of subtelomeric genes (**Figure 2E**) and short telomeres (**Figure 2F**), equivalent to complete loss of Set1. While these data may indicate that Set1 catalytic activity is the primary requirement to support telomere maintenance, the decreased stability of Set1 with these mutations leaves open the possibility that alternate, non-catalytic functions for Set1 may still be required. Furthermore, these data underscore the importance of evaluating protein expression of methyltransferases when using mutants to distinguish catalytic and non-catalytic roles.

**Figure 2.**
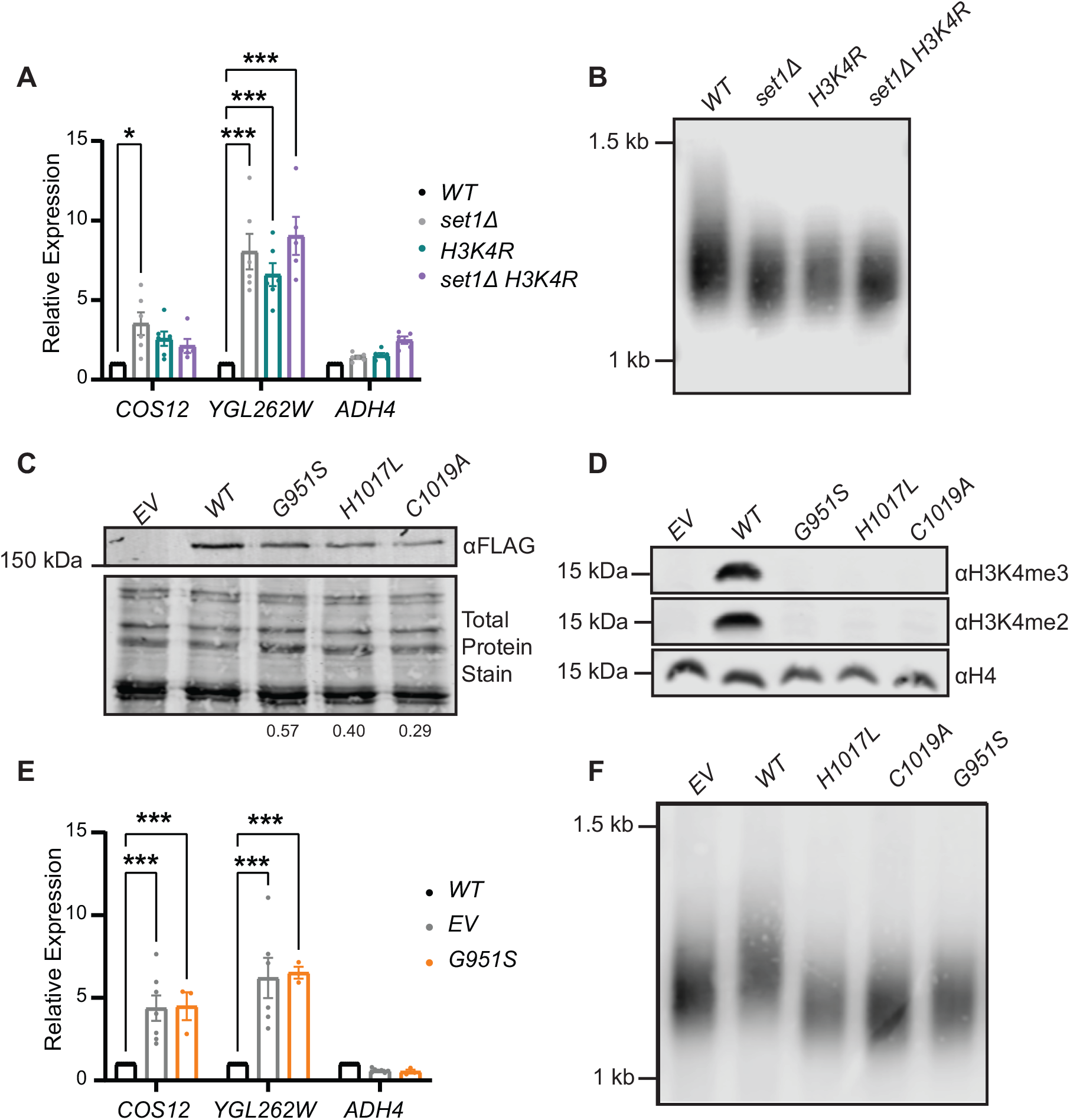
Analysis of catalytic requirements of Set1 in subtelomeric gene repression and telomere length control. (**A**) RT-qPCR analysis of *TEL07L* genes in *WT, set1Δ, H3K4R,* and *set1Δ H3K4R* cells. Expression was normalized to the control gene *TFC1* and is shown relative to *WT*. Error bars represent standard error of the mean (SEM) for a minimum of three biological replicates. (**B**) Southern blot showing terminal telomere fragment molecular weight in *WT, set1Δ, H3K4R*, and *set1Δ H3K4R* cells. (**C** and **D**) Western blots depicting FLAG-Set1 (**C**) protein levels and H3K4 methylation (**D**) in *set1Δ* cells expressing *FLAG-SET1* with catalytic mutations or an empty vector (EV). In panel **C**, the abundance of each Set1 mutant to wildtype was determined by image analysis relative to total protein stain (see *Methods)* and is shown at the bottom of the gel. (**E**) RT-qPCR analysis of *TEL07L* genes in *set1Δ* cells carrying an empty vector (EV) or expressing FLAG-Set1 *(WT)* or FLAG-Set1 G951S. Expression was normalized to the control gene *TFC1* and is shown relative to *WT.* Error bars represent standard error of the mean (SEM) for a minimum of three biological replicates. (**F**) Southern blot showing terminal telomere fragment molecular weight in *set1Δ* cells with empty vector (EV) or FLAG-Set1 catalytic mutants. For gene expression analysis, significance was evaluated using two-way ANOVA and Turkey’s multiple comparisons test. *p*-values are indicated as follows: * < 0.05, ** < 0.01, *** <0.001.

Set1 contains two RNA recognition motifs (RRM1 and RRM2) near the N-terminus, which predominantly bind mRNAs, as well as a small number of non-coding RNAs (60–62), and the catalytic core of Set1 is comprised of the N-SET, SET, and post-SET domains near the C-terminus of the protein (**Figure 3A**). To determine whether non-catalytic functions of Set1 contribute to its role at telomeres, we generated deletions of RRM1, RRM2, and a region spanning all of RRM1 and RRM2 (RRM1-2) in the *FLAG-SET1* construct (**Figure 3A**). Previous work (57,62) indicated that H3K4 methylation is moderately reduced in the absence of the RRM domains. Therefore, we also generated the reported hyperactive methylation mutant G990E (57) to help overcome this deficiency and potentially separate H3K4 methyltransferase activity from RNA binding activity. In addition, we tested the H422A point mutation, which is reported to block RNA binding by RRM2 yet maintain H3K4 methylation status. Immunoblotting of the FLAG-Set1 ΔRRM1, ΔRRM2, and ΔRRM1-2 proteins expressed from their endogenous promoter, as well as the H422A point mutant, showed similar expression to wildtype FLAG-Set1 for all mutant proteins (**Figure 3B** and **Supp. Figure 1A**). H3K4 methylation levels were also comparable to cells with FLAG-Set1, although some degree of H3K4me3 was lost in these mutants, even in combination with the *G990E* allele (**Figure 3C** and **Supp. Figure S1B**). This is likely due to the reported reduction in Set1 chromatin binding when the RRMs are absent (60).

**Figure 3.**
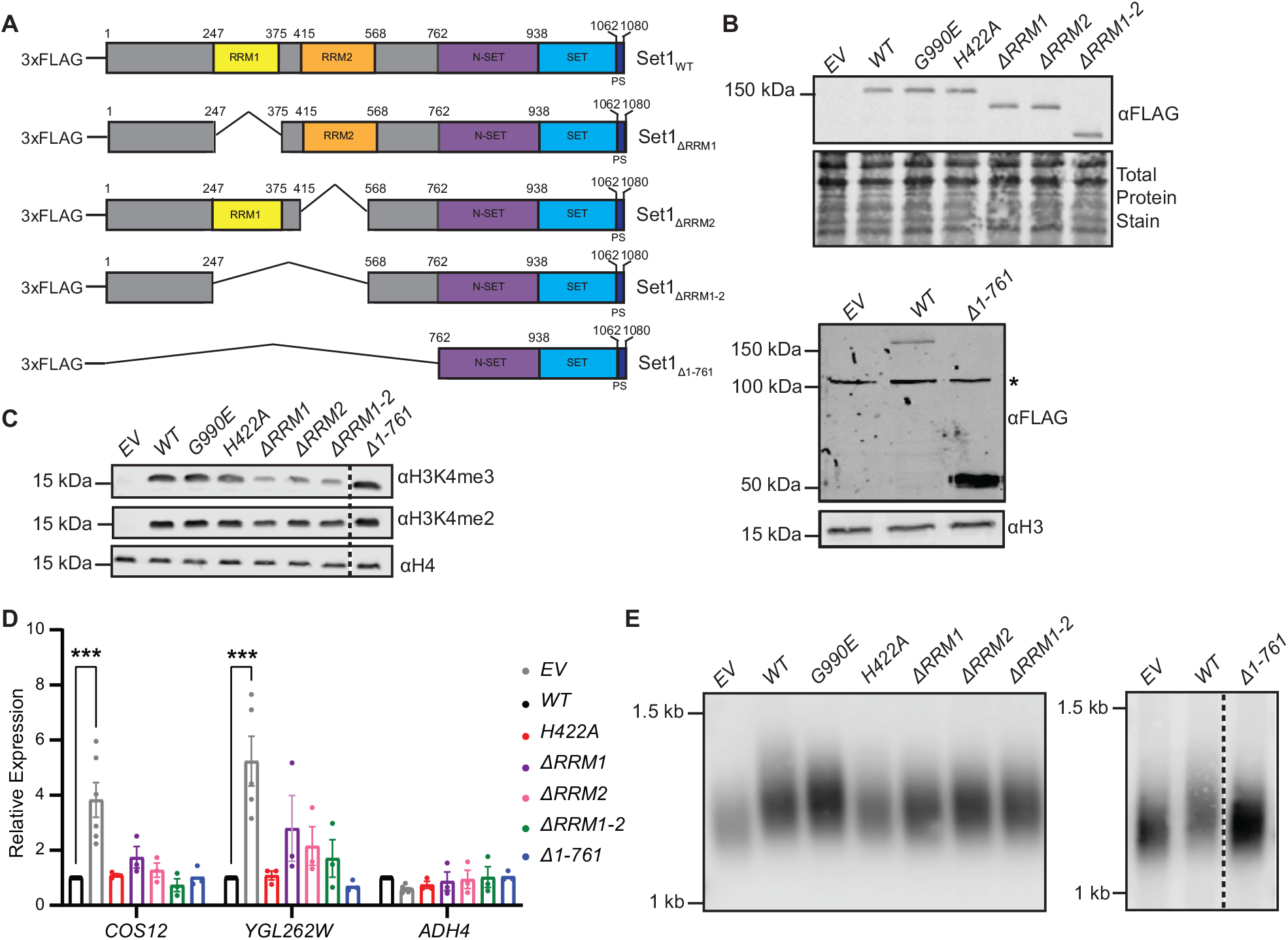
The catalytic core of Set1, but not the RRM domains, is required for subtelomeric gene repression and telomere length maintenance. (**A**) Schematic depicting wildtype Set1 domain structure (top) and domain deletions *(ΔRRM1, ΔRRM2, ΔRRM1-2*) and C-terminal truncation *(Δ1-761)* used in expression vectors with an N-terminal FLAG tag in *set1Δ* cells. (**B**) Western blots of yeast whole cell extracts probed with anti-FLAG showing levels of wildtype and mutant FLAG-Set1 expressed in *set1Δ* cells and (**C**) and H3K4me3 and H3K4me2 levels for each strain expressing FLAG-Set1 variants. Total protein stain, anti-H3, or anti-H4 were used as loading controls. Asterisk (*) indicates a non-specific band detected by the anti-FLAG antibody. (**D**) RT-qPCR analysis of *TEL07L* genes in *set1Δ* cells carrying an empty vector (EV) or expressing FLAG-Set1 *(WT)* or FLAG-Set1 mutants. Expression was normalized to the control gene *TFC1* and is shown relative to *WT*. Error bars represent standard error of the mean (SEM) for a minimum of three biological replicates. Significance was evaluated using two-way ANOVA and Turkey’s multiple comparisons test. *p*-values are indicated as follows: * < 0.05, ** < 0.01, *** <0.001. (**E**) Southern blot showing terminal telomere fragment molecular weight in *set1Δ* cells with empty vector (EV) or FLAG-Set1 mutants. In all gel image panels, the dashed line separates samples run on the same gel in which intervening lanes were removed for clarity.

Analysis of subtelomeric gene expression using RT-qPCR showed that the RNA binding activity of Set1 is dispensable for the maintenance of gene repression, as all mutants tested displayed repression of subtelomeric genes (**Figure 3D**). RNA-binding by Set1 also appears to be dispensable for maintaining telomere length, as the H422A mutant and RRM-deleted proteins (ΔRRM1, ΔRRM2, and ΔRRM1-2) displayed telomere lengths more similar to wildtype than *set1Δ* cells (**Figure 3E**). Similar assays using the FLAG-Set1 RRM-deleted protein combined with the hyperactive G990E mutation were performed, however, there was no change in phenotype in the presence of this mutation (**Supp. Figure S1C** and **S1D**).

We next tested whether FLAG-Set1 containing only the catalytic core, residues 762 to 1081 (Δ1-761) (63), was sufficient to promote subtelomeric gene repression and telomere length maintenance. As demonstrated in previous work (63), FLAG-Set1 Δ1-761 is stable, highly expressed, and maintains wildtype H3K4me levels (**Figure 3B** and **3C**). Cells expressing FLAG-Set1 Δ1-761 closely mimicked the subtelomeric gene expression of cells containing full-length FLAG-Set1 (**Figure 3D**) and displayed telomere length consistent with the presence of a wildtype copy of Set1 (**Figure 3E**), indicating that the catalytic core of Set1 alone is sufficient to maintain telomere length and subtelomeric gene repression. These results are consistent with previous work indicating that overexpression of the SET domain of Set1 rescues the telomere position effect phenotype of *set1Δ* cells (14,19).

Set1 has also been implicated in DNA damage response (DDR) pathways (19,64). To determine whether Set1’s role in these pathways is also primarily dependent on the SET domain, we tested growth of strains expressing mutant FLAG-Set1 in the presence of hydroxyurea (HU) and caffeine (**Supp. Figure S1E**). Cells lacking Set1 (empty vector) grew poorly in the presence of either HU or caffeine, whereas the expression of FLAG-Set1 was able to rescue this growth defect. The RNA binding mutants of FLAG-Set1 (ΔRRM1, ΔRRM2, ΔRRM1-2, and H422A) behave similar to wildtype, whereas the catalytic mutants H1017L, C1019A, and G951S were strongly inhibited in the presence of HU or caffeine (**Supp. Figure S1E**). Expression of the FLAG-Set1 Δ1-761 catalytic core exhibited growth similar to full length FLAG-Set1, indicating that the catalytic core is also sufficient for Set1’s role in the DDR pathway (65).

### Subtelomeric gene repression, but not telomere length maintenance, depends on H3K4 methylation

In addition to Set1, COMPASS complex subunits and the Rad6-Bre1 H2B ubiquitin ligase complex promote H3K4 methylation: loss of COMPASS subunits Spp1 and Sdc1 lead to a reduction in H3K4me3 and H3K4me2, respectively (66), and *rad6Δ* mutants have no H3K4me due to the requirement for H2BK123Ub (67). We and others have previously shown that subtelomeric gene repression of *TEL07L* adjacent genes *COS12* and *YGL262W* is reduced in *spp1Δ* and *sdc1Δ* mutants (**Figure 4A**; (1,12,13)). However, there is less derepression in these cells compared to those lacking Set1, indicating a partial requirement for these COMPASS complex components in subtelomeric gene repression. We also monitored subtelomeric gene expression in *rad6Δ* cells, which have no H3K4 methylation, and observed that this mutant closely mimics the derepression observed in *set1Δ* cells (**Figure 4A**). In Southern blots of terminal telomere fragments, both *spp1Δ* and *sdc1Δ* mutants showed little defect in telomere length compared to isogenic wildtype cells (**Figure 4B**). Similarly, a modest defect in telomere length was observed in the *rad6Δ* mutant (**Figure 4B**; (24,68)), despite the clear loss of subtelomeric gene repression in these cells. These data suggest that the defects in telomere length maintenance and subtelomeric gene expression can be separated in some mutants lacking H3K4 methylation. Furthermore, while our data indicate a requirement for the catalytic core of Set1 in these telomere maintenance phenotypes, there appears to be only a partial requirement for H3K4 methylation, particularly for telomere length regulation.

**Figure 4.**
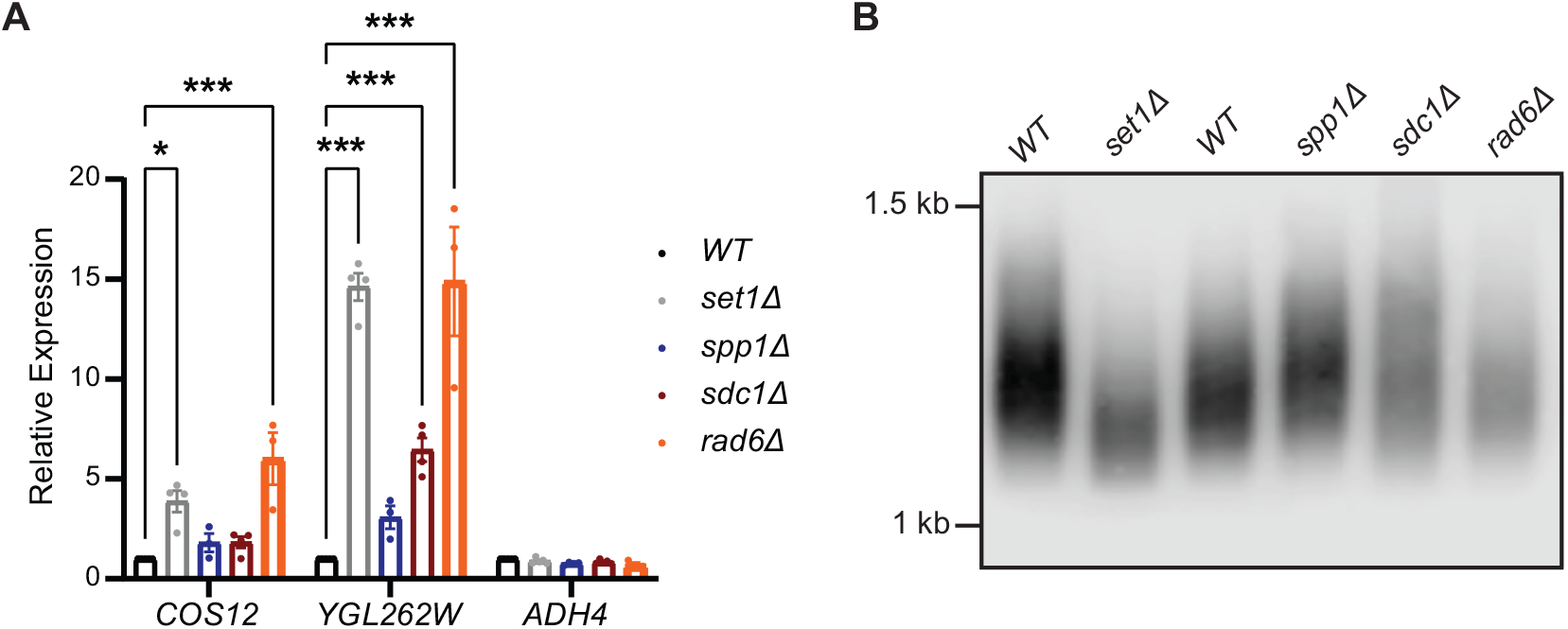
The requirement for other COMPASS components and Rad6 in subtelomeric gene repression and telomere length. (**A**) RT-qPCR analysis of *TEL07L* genes in the indicated deletion mutant strains. Expression was normalized to the control gene *TFC1* and is shown relative to *WT*. Error bars represent standard error of the mean (SEM) for a minimum of three biological replicates. Significance was evaluated using two-way ANOVA and Turkey’s multiple comparisons test. *p*-values are indicated as follows: * < 0.05, ** < 0.01, *** <0.001. (**B**) Southern blot showing terminal telomere fragment molecular weight in the indicated mutants.

### The abundance of telomere maintenance factors are altered upon loss of Set1

Next, we sought to better define the potential roles for Set1 in telomere length maintenance and subtelomeric gene repression. We and others have previously suggested that Set1 and H3K4 methylation likely support telomere maintenance through additional mechanisms beyond regulating repressive chromatin factors at subtelomeres (9,12). This observation is also supported by our finding that the transcriptome of *set1Δ* mutants is highly correlated with the transcriptome of yeast cells lacking components of the telomerase holoenzyme, such as *est1Δ, est3Δ,* and *tlc1Δ,* as well as mutants in other telomere maintenance pathways (12).

To determine whether any telomere maintenance factors are misregulated in the absence of Set1, we assayed mRNA levels of a subset of relevant genes in *set1Δ* cells using RT-qPCR (**Figure 5A**). In these mutants, members of the CST capping complex (Cdc13-Stn1-Ten1) showed altered steady state mRNA levels, with higher levels of *CDC13* and *STN1* mRNAs, and lower levels of *TEN1* mRNA. In addition, the mRNAs coding for telomerase subunits *EST1, EST2,* and the telomerase RNA, *TLC1* all showed increased abundance in the absence of Set1, whereas the *EST3* mRNA is two-fold less abundant. The mRNA abundance of additional telomere maintenance regulators, including those coding for the transcription factor Rap1 and members of the RFA, yKu, and Rif1/2 complexes were also tested, however, there were few statistically significant changes in mRNA levels in *set1Δ* cells compared to wildtype (**Figure 5A**).

**Figure 5.**
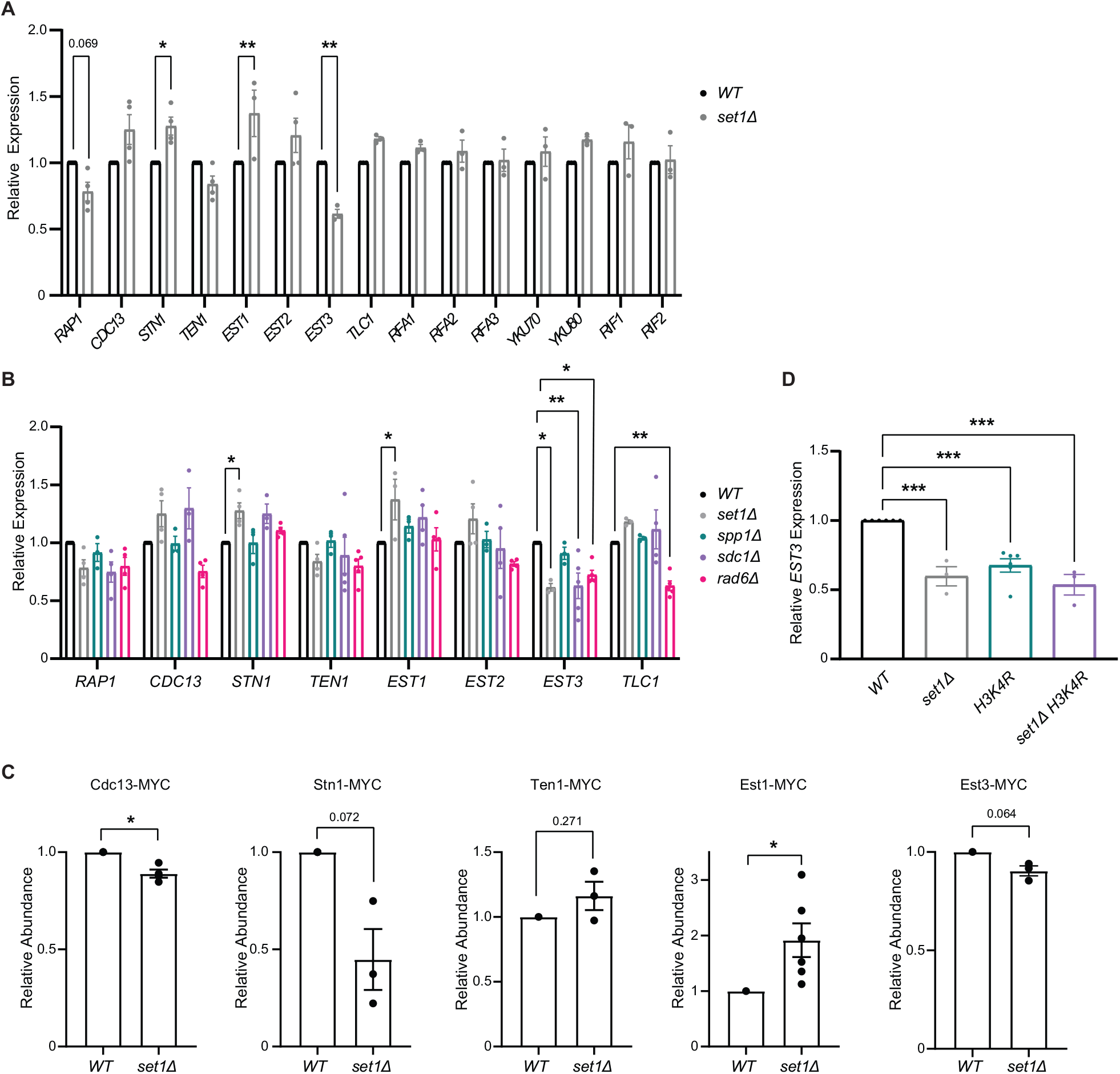
Genes encoding telomere maintenance factors show altered abundance levels in *set1Δ* cells. (**A** and **B**) RT-qPCR analysis of *TEL07L* genes in the indicated deletion mutant strains. Expression was normalized to the control gene *TFC1* and is shown relative to *WT*. Error bars represent standard error of the mean (SEM) for a minimum of three biological replicates. Significance was evaluated using two-way ANOVA and Sidak’s multiple comparisons test (**A**) or one-way ANOVA and Turkey’s multiple comparisons test for individual genes (**B**). (**C**) Quantification of protein abundances of Cdc13-MYC, Stn1-MYC, Ten1-MYC, Est1-MYC, and Est3-MYC in wildtype and *set1Δ* cells. Intensity of MYC signal was determined by Western blot and normalized using a total protein stain as a loading control for a minimum of three biological replicates. Error bars represent standard error of the mean (SEM). Significance was evaluated using an unpaired t-test. (**D**) RT-qPCR analysis of *EST3* in the indicated mutants. Data presented as described for panel **A.** For all panels, *p*-values are indicated as follows: * < 0.05, ** < 0.01, *** <0.001.

We also tested whether mRNA abundance of a subset of telomere maintenance factors is altered in the *spp1Δ, sdc1Δ,* and *rad6Δ* mutants (**Figure 5B**). While some changes in mRNA levels were observed in the *spp1Δ* cells lacking H3K4me3, the *sdc1Δ* mutant most closely resembled *set1Δ* cells, indicating that loss of H3K4me2 is likely a main factor in disrupting the levels of mRNAs encoding telomere maintenance proteins in *set1Δ* cells. The *rad6Δ* mutant showed the most difference from the profile observed in *set1Δ* cells. While this may in part be due to loss of H3K4 methylation, it also likely stems from other functions for Rad6, including its role in promoting H3K79 methylation (24,68,69).

Previous reports indicate that the expression levels and stoichiometry of telomere regulators such as CST and telomerase are highly calibrated by the cell (35–46). Our analysis of mRNA abundance indicates that the levels of these proteins are likely changing in the absence of Set1. To monitor protein abundance of the CST and telomerase complexes, we epitope tagged CST and telomerase subunits in wildtype and *set1Δ* cells and performed quantitative immunoblotting (**Figure 5C**). The target protein abundance was monitored using an anti-MYC antibody and total protein levels were determined by staining the blot with a general protein stain. Cdc13-Myc levels were slightly but significantly reduced in *set1Δ* relative to wildtype and Stn1-Myc protein levels appear to be more reduced in cells lacking Set1, whereas Ten1 levels do not significantly differ in *set1Δ* cells (**Figure 5C**). Interestingly, the altered abundance of Stn1 protein does not parallel the changes in *STN1* mRNA expression, suggesting that it may be subject to translational or post-translational control in the absence of Set1.

The telomerase subunit Est1 showed increased protein abundance in *set1Δ* cells, consistent with higher levels of *EST1* mRNA in the mutant (**Figure 5A** and **5C**). Intriguingly, our analysis of Est3-MYC protein levels showed only a minor decrease in Est3-MYC abundance in *set1Δ* cells compared to wildtype (**Figure 5C**). This was surprising, given that the *EST3* mRNA abundance was the most substantially decreased in the absence of Set1 (**Figure 5A**). It was also decreased in the *sdc1Δ* and *rad6Δ* mutants (**Figure 5B**), and the *H3K4R* mutant showed similarly reduced levels of *EST3* mRNA (**Figure 5D**), indicating that the abundance of this mRNA is likely dependent on functional Set1 and H3K4 methylation levels.

### *Regulation of* EST3 *mRNA in set1Δ mutants*

To better understand the relationship between *EST3* mRNA and protein levels, we also evaluated the steady-state levels of the mRNA encoding *EST3-MYC.* This mRNA had equivalent abundance in wildtype and *set1Δ* cells (**Figure 6A**), unlike the endogenous *EST3*, however its levels were substantially decreased compared to the endogenous mRNA (**Figure 6A**). Currently, immunoblotting of the endogenous protein is not feasible, as there is not an available antibody for the recognition of yeast Est3. Previous reports have indicated that epitope tags disrupt the function of Est3 (44,45); therefore, we evaluated the function of our Est3-MYC construct in subtelomeric gene repression and telomere length. Wildtype cells expressing Est3-MYC showed a decrease in subtelomeric gene repression compared with wildtype cells expressing the endogenous untagged Est3, an effect that was further exacerbated in combination with *set1Δ* (**Figure 6B**). We also observed that cells with Est3-MYC have shorter telomeres, approximately equivalent to *set1Δ* cells (**Figure 6C**). However, *set1Δ* mutants with *EST3-MYC* show even shorter telomeres than *set1Δ* alone, indicating that each allele is contributing to the short telomere phenotype through distinct pathways. These data suggest that the *EST3-MYC* allele causes a loss-of-function phenotype due to decreased mRNA abundance and disrupted protein function, and it is not likely to accurately reflect endogenous protein levels of Est3.

**Figure 6.**
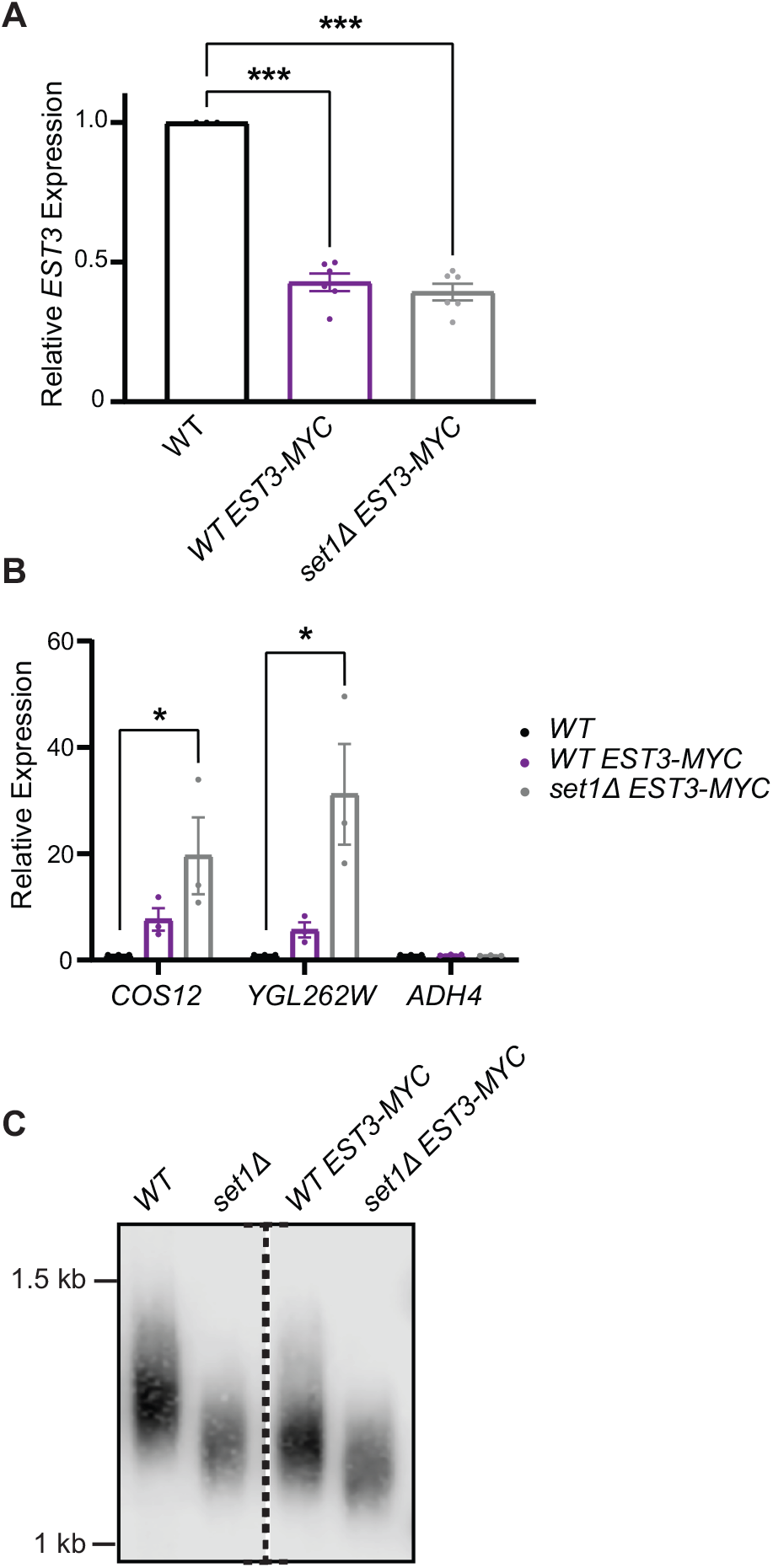
Incorporation of a C-terminal MYC tag disrupts *EST3* mRNA abundance, subtelomeric gene repression, and telomere length. (**A**) RT-qPCR analysis of *EST3-MYC* in *WT* and *set1Δ* cells carrying an integrated allele. (**B**) RT-qPCR analysis of *TEL07L* genes in *EST3-MYC* strains. For **A** and **B**, expression was normalized to the control gene *TFC1* and is shown relative to *WT*. Error bars represent standard error of the mean (SEM) for a minimum of three biological replicates. Significance was evaluated using one-way ANOVA and Dunnett’s multiple comparisons test (**A**) or two-way ANOVA and Sidak’s multiple comparisons test (**B**). *p*-values are indicated as follows: * < 0.05, ** < 0.01, *** <0.001. (**C**) Southern blot showing terminal telomere fragment molecular weight in the indicated strains. The dashed line in the blot indicates samples run on the same gel in which intervening lanes were removed for clarity.

Despite the complications in accurately evaluating Est3 protein levels, we next addressed whether Set1 may regulate the transcription of *EST3.* Given that other mutants that show reduced H3K4 methylation also showed lower levels of *EST3* mRNA (**Figure 5**), we performed chIP of H3K4me3 and H3K4me2 at the *EST3* promoter, 5’ ORF and 3’ ORF. We compared this to levels at similar regions of *EST1,* for which the mRNA showed increased abundance in the absence of Set1. Both H3K4me3 and H3K4me2 were present to a similar extent in the promoters and 5’ ends of *EST3* and *EST1,* however there was a marked difference at the 3’ end of the ORFs (**Figure 7A** and **7B**). Both histone marks, though particularly H3K4me2, showed increased levels at the 3’ end of *EST3* compared to *EST1,* which may suggest a greater requirement for H3K4 methylation in the regulation of *EST3.*

**Figure 7.**
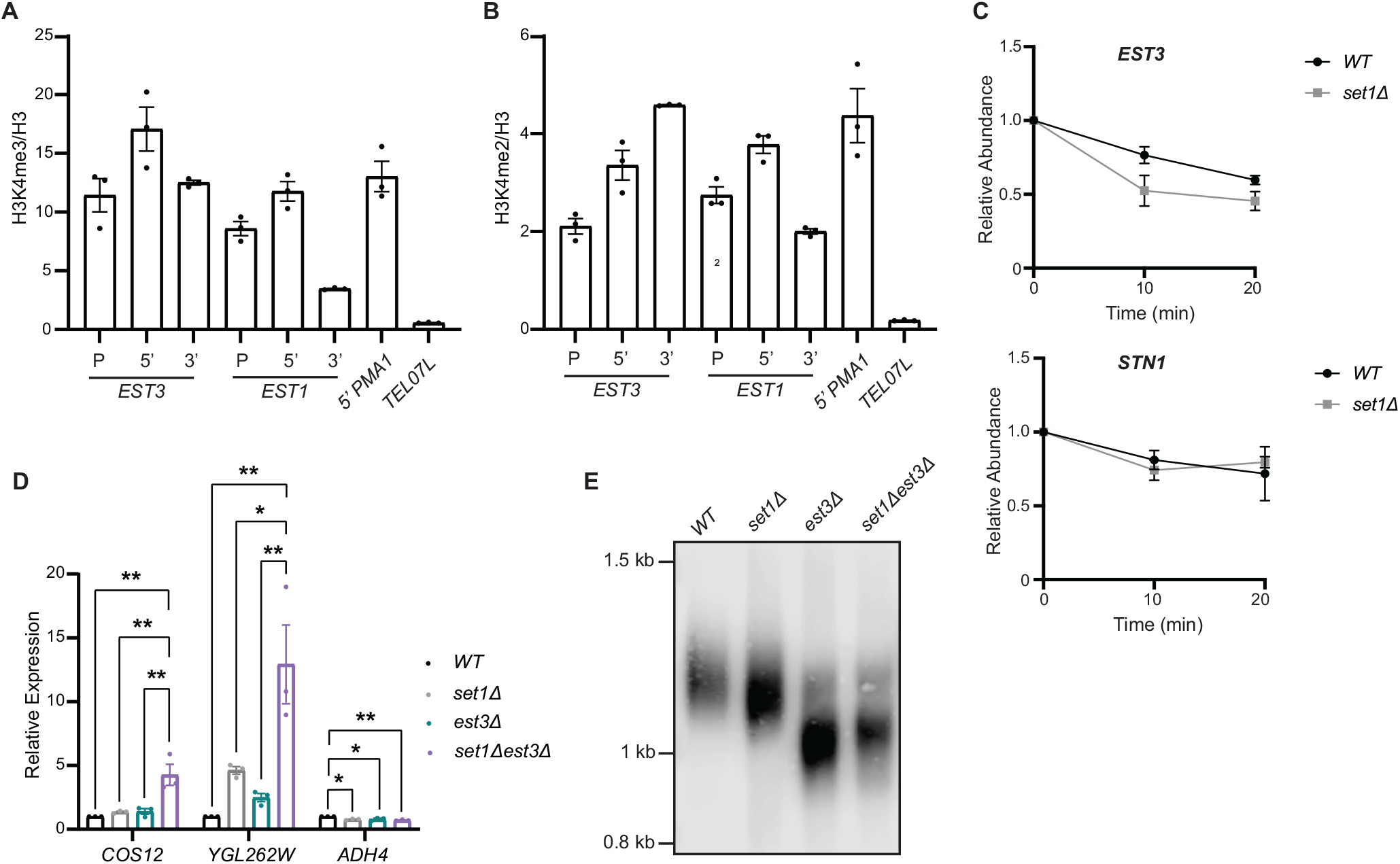
Regulation of endogenous *EST3* mRNA and epistasis analysis of *set1Δ est3Δ* mutants. Chromatin immunoprecipitation of H3K4me3 (**A**) and H3K4me2 (**B**) shown relative to total H3 at *EST3* and *EST1* gene loci. Primer sets amplify promoter (P), 5’ coding region (5’), and 3’ coding regions (3’). The 5’ coding region of *PMA1* serves as a positive control for H3K4me3/H3K4me2 binding, and the *TEL07L* region is a negative control. SEM of three biological replicates is shown. (**C**) Relative abundance of *EST3* and *STN1* mRNAs in *WT* and *set1Δ* cells at 10 and 20 minutes after thiolutin treatment. *EST3* and *STN1* levels were normalized to the RNA pol II-independent transcript *sCR1.* SEM of three biological replicates are shown. (**D**) RT-qPCR analysis of *TEL07L* genes in the indicated deletion mutant strains. Expression was normalized to the control gene *TFC1* and is shown relative to *WT*. Error bars represent standard error of the mean (SEM) for a minimum of three biological replicates. Significance was evaluated using two-way ANOVA and Turkey’s multiple comparisons test. *p*-values are indicated as follows: * < 0.05, ** < 0.01, *** <0.001. (**E**) Southern blot showing terminal telomere fragment molecular weight in the indicated strains.

We also evaluated whether Set1 may regulate post-transcriptional processing or stability of the *EST3* mRNA. In this assay, we compared *EST3* abundance to that of *STN1* since its protein levels appeared decreased by immunoblot despite increased abundance of its mRNA (**Figure 5B** and **C**). Wildtype and *set1Δ* cells were treated with the transcription inhibitor thiolutin and mRNA abundance of each transcript was measured by RT-qPCR and compared to the RNA pol III-controlled transcript *sCR1.* In this assay, we observed more rapid turnover of the *EST3* transcript in *set1Δ* cells compared to wildtype (**Figure 7C**; *p-*value = 0.013 from two-way ANOVA), indicating a shorter half-life for the *EST3* mRNA upon loss of Set1. This differed from *STN1,* which showed similar levels in both wildtype and *set1Δ* cells following thiolutin treatment (*p-*value = 0.976 from two-way ANOVA), suggesting each of these transcripts is subject to different regulatory mechanisms dependent on Set1.

### Overlapping functions of Set1 and Est3 in telomere maintenance phenotypes

To investigate the dependence of telomere maintenance defects in *set1Δ* on *EST3* expression, we generated double *set1Δ est3Δ* mutants following sporulation of heterozygous diploids and evaluated subtelomeric gene repression and telomere length. As expected, both *set1Δ* and *est3Δ* single mutants show increased subtelomeric gene expression (**Figure 7D**), however there is a synergistic increase in expression in *set1Δ est3Δ* double mutants. These data suggest that both Set1 and Est3 are separately required for subtelomeric gene repression, and loss of each of their activities disrupts multiple pathways normally repressing genes within subtelomeric chromatin. Both *set1Δ* and *est3Δ* single mutants also exhibited shorter telomeres (**Figure 7E**), however *est3Δ* mutants were much shorter than *set1Δ* mutants. Interestingly, the *set1Δ est3Δ* double mutants showed similar telomere length to *est3Δ* single mutants, indicating that Set1 and Est3 may contribute to the same pathway required for maintaining appropriate telomere lengths. These data argue that one potential factor contributing to short telomeres in *set1Δ* cells is reduced *EST3* mRNA abundance, however this is not likely a key contributor to the misregulation of subtelomeric gene repression in the absence of Set1.

## Discussion

The lysine methyltransferase Set1 has long been implicated in the regulation of subtelomeric gene repression and telomere length maintenance (2,12,14,19–21), yet its specific role in these processes has not been completely defined. Based on the association of Set1 and H3K4 methylation with active transcription, the euchromatic distribution of H3K4me3 has been implicated in the maintenance of repressive chromatin at subtelomeres (10,22). However, other observations suggest that there are likely additional mechanisms that underlie Set1’s role in telomere maintenance pathways. For example, loss of COMPASS components and other factors that deplete H3K4 methyl levels do not always display phenotypes similar to *set1Δ* cells (24,68). In addition, we previously reported that the transcriptome profile of *set1Δ* mutants is most highly correlated with mutants in genes directly linked to telomere maintenance, such as those encoding telomerase subunits (12). Given these observations, we sought to more systematically define the role for Set1 and H3K4 methylation in two metrics of telomere health: chromatin-mediated gene repression and telomere length.

### The Set1 SET domain is required for telomere health through H3K4-methylation dependent and independent pathways

Previous studies using reporter assays have suggested that the SET domain of Set1 is primarily responsible for maintaining subtelomeric gene repression (14,19), however the specific contribution of Set1 catalytic methyltransferase activity at H3K4 has been difficult to assess. We used a series of genetic mutations, including the H3K4R point mutation, COMPASS mutants, and mutations in Set1, to quantitatively compare subtelomeric gene expression and simultaneously evaluate telomere length. In a *set1Δ* background, we expressed *FLAG-SET1* from its endogenous promoter on a plasmid and generated multiple point mutations known to abrogate catalytic activity. As loss of catalytic activity is reported to destabilize the Set1 protein (56), we monitored abundance of these mutants to clarify whether or not these specific mutations promote Set1 instability. In all mutants tested, we observed decreased levels of FLAG-Set1, indicating that use of these mutants does not provide clean separation of Set1 catalytic and non-catalytic functions.

In addition to catalytic activity localized within the SET domain, Set1 contains two RNA recognition motifs (RRMs), which bind a diversity of RNAs (57,60,61), though whether they contribute to Set1-dependent telomere maintenance had not been directly investigated. Previous studies have also differed on the extent to which H3K4 methylation is affected by loss of the RRM domains (57,62). However, our immunoblotting shows equivalent expression of all Set1 mutants tested with moderate loss of H3K4me3 in the RRM deletion mutants, although not in the H422A point mutation that disrupts RNA binding (62). This mutant analysis showed that the catalytic core of Set1 is predominantly responsible for maintaining subtelomeric gene repression and telomere length. Interestingly, while the expression of the Set1 catalytic core (Set1 Δ1-761) restores H3K4 methylation levels, the distribution of H3K4me3 and H3K4me2 in the genome is disrupted (70). This suggests that specific placement of the H3K4 methyl marks is not required for Set1-dependent telomere maintenance, or that an alternate function of the catalytic core of Set1 may be required. In addition, our analysis of gene expression and telomere length in COMPASS mutants *spp1Δ* and *sdc1Δ,* as well as *rad6Δ,* showed that subtelomeric gene expression changes correlated with the relative loss of H3K4 methyl species in these mutants. However, this was not the case for telomere length, which showed little change from wildtype in all of these mutants. These findings indicate that telomere length maintenance and repression of subtelomeric chromatin are separable processes that show differential requirements for H3K4 methylation.

In addition to catalyzing H3K4 methylation, the SET domain of Set1 has additional functions. Set1 methylates the kinetochore-associated protein Dam1 (26), and more recently, H3K37me1 has been reported to be targeted by Set1 together with the H3K36 methyltransferase Set2 (27). In a non-catalytic role, the SET domain of Set1 physically interacts with the DNA damage protein Mec3 (19), which may also contribute to its function at telomeres. Furthermore, orthologs of Set1 have been shown to perform functions independent of the catalytic methyltransferase activity (28–31). Further identification of mutants that separate functions of the SET domain will be needed to better define the specific contributions of catalytic and non-catalytic activities to Set1’s role in telomere maintenance, as well as its other biological roles.

### Altered abundance of telomere maintenance factors in set1Δ mutants through transcriptional and post-transcriptional regulation

To better define the role of Set1 in telomere maintenance, we assayed steadystate mRNA and protein abundance of a series of telomere regulators in cells lacking Set1, Spp1, Sdc1, or Rad6, with a focus on components of the CST telomere capping complex and the telomerase holoenzyme. At the mRNA level, we observed alterations to the abundance of *STN1, TEN1, EST1, EST3,* and *TLC1.* There appeared to be an enhanced dependence on H3K4me2 or H3K4me1 for regulating mRNA abundance, as *sdc1Δ* mutants most closely mimicked *set1Δ* cells. Interestingly, in monitoring protein abundance of these factors, we observed that the mRNA and protein levels change in parallel for some factors, such as Est1, however some proteins displayed changes in abundance that could not be explained by changes at the mRNA level. For example, *STN1* transcript levels are increased in *set1Δ,* while Stn1-MYC was decreased. This is also seen, albeit to a lesser extent, with Cdc13, whose transcript levels increased in *set1Δ* cells while the protein abundance is slightly reduced. This suggests altered translational or post-translational regulation of CST components in the absence of Set1. Both Cdc13 and Stn1 are subject to post-translational regulation: they are phosphorylated by the cyclin-dependent kinase, Cdk1, and this modification serves to stabilize the complex at the telomere (37,38,71). This phosphorylation, and possibly other regulatory mechanisms including transcriptional control, may be disrupted in *set1Δ* cells due to their aberrant cell cycle progression (72). The abundance and stoichiometry of CST complex components is exquisitely regulated in the cell, and previous work has identified multiple mechanisms of regulation and shown that alterations to any of the complex members can impact telomere function and promote feedback regulation of other telomere maintenance factors (35–46). Our data indicate that Set1 promotes the proper balance of CST components within the cell and this disruption in *set1Δ* cells likely contributes to impaired telomere function.

In addition to altered abundance of CST factors, we noted differential abundance of *EST1* and *EST3* mRNAs, each of which encode components of the telomerase holoenzyme. The increase in *EST1* in *set1Δ* cells was reflected in higher abundance of the Est1-MYC protein. At the transcript level, *EST3* showed the largest change in abundance in *set1Δ* cells and related mutants. This is consistent with our previous findings that the transcriptome in *set1Δ* cells is highly correlated with that in *est3Δ* mutants (12). Further analysis of *EST3* mRNA half-life following thiolutin treatment indicated more rapid degradation of *EST3* mRNA than that of *STN1,* suggesting that *EST3* post-transcriptional regulation is disrupted in the absence of Set1. Interestingly, the *EST3* mRNA undergoes programmed ribosomal frameshifting and is regulated by the nonsense mediated decay (NMD) pathway (41,46). One possibility is that there is increased shunting of the *EST3* mRNA to the NMD pathway in *set1Δ* cells, potentially through misregulated frameshifting or other processing or translation defect.

We investigated Est3 protein abundance in *set1Δ* cells by integrating a MYC tag at the C-terminus, as there is no available antibody for detecting Est3. It has previously been reported that C-terminal epitope tags can disrupt function of Est3 (44,45). Indeed, we observed decreased gene repression and shorter telomeres with the addition of the C-terminal MYC tag, though also a substantial decrease in mRNA abundance. While we could not conclusively determine whether full-length Est3 protein showed decreased abundance in *set1Δ* cells, epistasis analysis of *set1Δ est3Δ* double mutants suggested that Set1 and Est3 contribute separately to subtelomeric gene repression. However, *set1Δ est3Δ* cells have similar length telomeres to *est3Δ* single mutants, indicating that Set1 and Est3 both contribute to the same pathway to maintain telomere length.

In summary, our data indicate that Set1’s role at telomeres depends on the activity of its catalytic core, although H3K4 methylation-independent functions likely contribute to telomere length maintenance. The role for Set1 in subtelomeric gene repression is most closely correlated with H3K4 methyl status, indicating this mark is required for gene repression, and, further, that misregulated subtelomeric chromatin is not the primary driver of telomere shortening in *set1Δ* mutants. These differential requirements for H3K4 methylation are also evident in our analysis of mRNA abundance of telomere maintenance factors, with genes such as *EST3* likely dependent on H3K4 methyl status for regulation, but not *EST1* or *STN1.* These transcripts show disrupted transcriptional and post-transcriptional regulation in the absence of Set1, as well as possible translational or post-translational mechanisms directing protein abundance. This likely reflects the multiple mechanisms which precisely calibrate the abundance of these critical telomere regulators. Altogether, this study provides new and unexpected insights into how Set1 promotes telomeric and subtelomeric function and provides the foundation for dissecting the roles of H3K4 methyl species and additional targets of Set1 in telomere maintenance pathways in yeast and other systems.

## Supporting Information

This article contains supporting information.

## Acknowledgements

The authors thank all members of the Green lab for helpful discussions, technical assistance, and feedback on the manuscript. We thank Dr. David Lydall for yeast strains and Dr. Philip Farabaugh for sharing equipment.

## Funding and Additional Information

This work was support by the National Institutes of Health R01GM124342 to EMG. MYN was supported by funds from NIH T32GM055036 and R25GM066706 and NSF LSAMP BD 1500511 to UMBC. The content is solely the responsibility of the authors and does not necessarily represent the official views of the National Institutes of Health.

