## Supplemental Tables and Figures for "Set1 regulates telomere function via H3K4 methylation-dependent and independent pathways and calibrates the abundance of telomere maintenance factors"

**Supplemental Figures**

**Figure S1**. Telomere-related phenotypes in *SET1* G990E mutants and RRM deletions. (**A**) Western blots of yeast whole cell extracts probed with anti-FLAG showing levels of wildtype and mutant FLAG-Set1 and (**B**) and H3K4me3 and H3K4me2 levels for each strain expressing FLAG-Set1 variants. Total protein stain or anti-H4 were used as loading controls. (**C**) RT-qPCR analysis of *TEL07L* genes in *set1Δ* cells carrying an empty vector (EV) or expressing FLAG-Set1 (*WT*) or FLAG-Set1 mutants. Expression was normalized to the control gene *TFC1* and is shown relative to *WT*. Error bars represent standard error of the mean (SEM) for a minimum of three biological replicates. Significance was evaluated using two-way ANOVA and Turkey’s multiple comparisons test. *p*-values are indicated as follows: * < 0.05, ** < 0.01, *** <0.001. (**D**) Southern blot showing terminal telomere fragment molecular weight in *set1Δ* cells with empty vector (EV) or FLAG-Set1 mutants. (**E**) Ten-fold serial dilutions of saturated yeast cultures were spotted on SC-URA, SC-URA + 100 mM HU, and SC-URA +10 mM caffeine plates and grown at 30°C.

**Supplemental Tables**

**Table S1. Yeast strains used in this study.**

| **Strain Number** | **MAT** | **Genotype** | **Reference** |
| --- | --- | --- | --- |
| yEG001 | a | *his3Δ1 leu2Δ0 met15Δ0 ura3Δ0 (WT)* | BY4741 |
| yEG230 | alpha | *his3Δ1 leu2Δ0 met15Δ0 ura3Δ0 (WT)* | (1) |
| yEG232 | a | *set1Δ::KANMX* | (1) |
| yEG1108 | alpha | *set1Δ::NATMX* | This study |
| yEG323 | alpha | *tlc1Δ::NATMX* | This study |
| yEG880 | a/alpha | *TLC1/tlc1Δ::NATMX SET1/set1Δ::KANMX* | This study |
| DLY3001 | a | *W303 RAD5^+^(WT)* | (2) |
| yEG1296 | alpha | *W303 RAD5^+^set1Δ::NATMX* | This study |
| yEG1297 | alpha | *W303 RAD5^+^ cdc13-1 int* | (2) |
| yEG1298 | a | *W303 RAD5^+^ set1Δ::NATMX cdc13-1 int* | This study |
| yEG108 | a | *MAT a, ura3–52, lys2–801, ade2–101, trp1Δ63, his3Δ200, leu2Δ1, hht1-hhtf1∷LEU2, hht2-hht2∷HIS), YCp50-copyII (ura3+,HHT2-HHF2) pRS314-HHT2-HHF2* | (3) |
| yEG109 | a | *MAT a, ura3–52, lys2–801, ade2–101, trp1Δ63, his3Δ200, leu2Δ1, hht1-hhtf1∷LEU2, hht2-hht2∷HIS), YCp50-copyII (ura3+,HHT2-HHF2) pRS314-HHT2 K4R-HHF2* | (3) |
| yEG1342 | a | *MAT a, ura3–52, lys2–801, ade2–101, trp1Δ63, his3Δ200, leu2Δ1, hht1-hhtf1∷LEU2, hht2-hht2∷HIS), YCp50-copyII (ura3+,HHT2-HHF2) pRS314-HHT2-HHF2 set1Δ::KANMX* | This study |
| yEG1343 | a | *MAT a, ura3–52, lys2–801, ade2–101, trp1Δ63, his3Δ200, leu2Δ1, hht1-hhtf1∷LEU2, hht2-hht2∷HIS), YCp50-copyII (ura3+,HHT2-HHF2) pRS314-HHT2 K4R-HHF2 set1Δ::KANMX* | This study |
| yEG100 | a | *spp1Δ::KANMX* | (1) |
| yEG110 | a | *sdc1Δ::KANMX* | (1) |
| yEG623 | alpha | *rad6Δ::HIS3MX* | (4) |
| yEG647 | a | *set1Δ::KANMX + p045 (pRS316)* | This study |
| yEG738 | a | *set1Δ::KANMX + p319 (pRS316FLAG-SET1)* | This study |
| yEG740 | a | *set1Δ::KANMX + p342 (pRS316 FLAG-SET1 G990E)* | This study |
| yEG741 | a | *set1Δ::KANMX + p343 (pRS316 FLAG-SET1 H422A)* | This study |
| yEG746 | a | *set1Δ::KANMX + p344 (pRS316 FLAG-SET1 ΔRRM1)* | This study |
| yEG747 | a | *set1Δ::KANMX + p345 (pRS316 FLAG-SET1 ΔRRM2)* | This study |
| yEG748 | a | *set1Δ::KANMX + p346 (pRS316 FLAG-SET1 ΔRRM1- ΔRRM2)* | This study |
| yEG819 | a | *set1Δ::KANMX + p363 (FLAG-SET1 G990E ΔRRM1)* | This study |
| yEG820 | a | *set1Δ::KANMX + p364 (FLAG-SET1 G990E ΔRRM2)* | This study |
| yEG821 | a | *set1Δ::KANMX + p365 (FLAG-SET1 G990E ΔRRM1- ΔRRM2)* | This study |
| yEG885 | a | *set1Δ::KANMX + p392 (pRS316-FLAG-SET1 H1017L)* | This study |
| yEG886 | a | *set1Δ::KANMX + p393 (pRS316 FLAG-SET1 C1019A)* | This study |
| yEG984 | a | *set1Δ::KANMX + p457 (pRS316 FLAG-SET1 G951S)* | This study |
| yEG1102 | a | *set1Δ::KANMX + p510 (pRS316 FLAG-SET1Δ1-761)* | This study |
| yEG1137 | a | *cdc13::CDC13-MYC::HIS3MX* | This study |
| yEG1138 | a | *set1Δ::KANMX cdc13::CDC13-MYC::HIS3MX* | This study |
| yEG1284 | alpha | *ten1::TEN1-MYC::HIS3MX* | This study |
| yEG1285 | a | *set1Δ::KANMX ten1::TEN1-MYC::HIS3MX* | This study |
| yEG1008 | alpha | *est1::EST1-MYC::HIS3MX* | This study |
| yEG1009 | alpha | *set1Δ::KANMX est1::EST1-MYC::HIS3MX* | This study |
| DLY5761 | alpha | *W303 RAD5^+^ stn1::STN1-MYC::TRP1* | (2) |
| yEG1318 | a | *W303 RAD5^+^ stn1::STN1-MYC::TRP1 set1Δ::NATMX* | This study |
| yEG1358 | a | *est3::EST3-MYC::HIS3MX* | This study |
| yEG1359 | a | *set1Δ::KANMX est3::EST3-MYC::HIS3MX* | This study |
| yEG1364 | a | *est3Δ::HygB* | This study |
| yEG1365 | a | *est3Δ::HygB set1Δ::KANMX* | This study |

Note: All strains used in this study are derived from the BY4741/BY4742 background unless otherwise indicated in the genotype column.

**Table S2. Plasmids used in this study.**

| **Plasmid Number** | **Description** | **Reference** |
| --- | --- | --- |
| p045 | pRS316 (URA3) |  |
| p340 | pRS316 + FLAG-SET1 | This study |
| p342 | pRS316 + FLAG-SET1 G990E | This study |
| p343 | pRS316 + FLAG-SET1 H422A | This study |
| p344 | pRS316 + FLAG-SET1 ΔRRM1 | This study |
| p345 | pRS316 + FLAG-SET1 ΔRRM2 | This study |
| p346 | pRS316 + FLAG-SET1 ΔRRM1- ΔRRM2 | This study |
| p363 | pRS316 + FLAG-SET1 ΔRRM1 G990E | This study |
| p364 | pRS316 + FLAG-SET1 ΔRRM2 G990E | This study |
| p365 | pRS316 + FLAG-SET1 ΔRRM1- ΔRRM2 G990E | This study |
| p392 | pRS316 + FLAG-SET1 H1017L | This study |
| p393 | pRS316 + FLAG-SET1 C1019A | This study |
| p510 | pRS316 + FLAG-SET1 Δ1-761 | This study |
| p457 | pRS316 + FLAG-SET1 G951S | This study |

**Table S3. Primers used in this study.**

| **Oligo Number** | **Experiment** | **ORF/Position** | **Sequence** |
| --- | --- | --- | --- |
| oEG537 | SYBR GEX | COS12 - F | CATTACAAATACTCCGGGTATAGACA |
| oEG538 | SYBR GEX | COS12 - R | GCAGCTGGAA CCATCAAAA |
| oEG539 | SYBR GEX | YGL262W - F | GAGAATTACTCTGACATTGGAGATGA |
| oEG540 | SYBR GEX | YGL262W - R | TTGTCATTAC AGAAGCCATC AAC |
| oEG541 | SYBR GEX | ADH4 - F | CCAATGTCACAGCTGGTTTG |
| oEG542 | SYBR GEX | ADH4 - R | CCTTAGCATTGTCGTGAGCA |
| oEG543 | SYBR GEX | TFC1 - F | ACACTCCAGGCGGTATTGAC |
| oEG544 | SYBR GEX | TFC1 - R | CTTCTGCAATGTTTGGCTCA |
| oEG1176 | SYBR GEX | TLC1 - F | TGTAGAAATCGCGCGTACTG |
| oEG1177 | SYBR GEX | TLC1 - R | CTATCCGCCTATCCTCGTCA |
| oEG1178 | SYBR GEX | RIF1 - F | AGTTCGTTGGCTGTTGAAGG |
| oEG1179 | SYBR GEX | RIF1 - R | TCGCTATCAGACGCATTTTG |
| oEG1351 | SYBR GEX | CDC13 - F | AAGAGCCTGAGTGTCCTCCA |
| oEG1352 | SYBR GEX | CDC13 - R | AATTGCACGGGAACTATTGC |
| oEG1353 | SYBR GEX | RAP1 - F | TAGCAACGTCAACGACGAAG |
| oEG1354 | SYBR GEX | RAP1 - R | GAAAAGATGCATTCCCCTCA |
| oEG1355 | SYBR GEX | EST2 - F | GTCACTTCAATGGCCTCGAT |
| oEG1356 | SYBR GEX | EST2 - R | CAGGAAGGCATGGTAATGCT |
| oEG1357 | SYBR GEX | STN1 - F | GCAAGAAGAACGCTTGAAGG |
| oEG1358 | SYBR GEX | STN1 - R | ACCGAAATGACAAGGAATGC |
| oEG1359 | SYBR GEX | TEN1 - F | TTCGCATTGGGGTATGGTAT |
| oEG1360 | SYBR GEX | TEN1 - R | CACGAACGTCATTCCTGGAT |
| oEG1361 | SYBR GEX | RIF2 - F | AAGTGTTAGCAGCCCAAAGG |
| oEG1362 | SYBR GEX | RIF2 - R | CTCTTGCAAGGCCATTGATT |
| oEG1417 | SYBR GEX | EST1 - F | GAACAACAGCGCAAAAGAGC |
| oEG1418 | SYBR GEX | EST1 - R | TCTGGGAGCGGAGACAATTT |
| oEG1419 | SYBR GEX | EST3 - F | TGAAGACAACTCGGAGCATG |
| oEG1420 | SYBR GEX | EST3 - R | AATTGTCGGGCTCATATGCG |
| oEG1421 | SYBR GEX | RFA1 - F | AGGGCTGGGAAGAAATTCGA |
| oEG1422 | SYBR GEX | RFA1 - R | GCTTGCTGATTCCATAGGCC |
| oEG1423 | SYBR GEX | RFA2 - F | CGGTGGCTTTGAGAACTCTG |
| oEG1424 | SYBR GEX | RFA2 - R | ATCGTCACAGGTGTCAAGGT |
| oEG1425 | SYBR GEX | RFA3 - F | CAATGATGACGGCGAGCTAG |
| oEG1426 | SYBR GEX | RFA3 - R | CTGTAAAGCAACCACACCGT |
| oEG1427 | SYBR GEX | yKU70 - F | GAGATTCCGGGTCAAAAGCA |
| oEG1428 | SYBR GEX | yKU70 - R | TATCTTGCGCCTCTTGTGGT |
| oEG1429 | SYBR GEX | yKU80 - F | TTCCCGTGACCATCTCCAAA |
| oEG1430 | SYBR GEX | yKU80 - R | TGATCGACTAGAACGGACGG |
| oEG1229 | Southern | telomeric TG(1-3) repeats | biotin-CACACCCACACCCACACC |
| oEG1637 | ChIP | EST1 promoter-F | CACAGACGAAGGTGCTTTCA |
| oEG1638 | ChIP | EST1 promote- R | TGAACGCGAAAATCACATTGA |
| oEG1639 | ChIP | EST1 5'-F | GCTCGTGCGCATCTGGATAA |
| oEG1640 | ChIP | EST1 5'-R | AGGAAGCATCTGAACGTGATAT |
| oEG1641 | ChIP | EST1 3'-F | AAACATGCTGCTTCACGAGG |
| oEG1642 | ChIP | EST1 3'-R | ATTGCTTCGTCTGGATATGAGAG |
| oEG1649 | ChIP | EST3 promoter-F | GTTCAATTCCCCGTCGCG |
| oEG1650 | ChIP | EST3 promoter-R | TGGTCAACTTTTGCTGTCTAGT |
| oEG1651 | ChIP | EST3 5'-F | TCATCCCTCTGGCCATGTAA |
| oEG1652 | ChIP | EST3 5'-R | CGAAATGGCACGGATTGGTT |
| oEG1653 | ChIP | EST3 3'-F | TTGGCGATGCTGACTTAGTC |
| oEG1654 | ChIP | EST3 3'-R | ACGGGCACTATTTCTTTGGAC |
| oEG153 | ChIP | 5'PMA1-F | TCAGCTCATCAGCCAACTCAAG |
| oEG154 | ChIP | 5'PMA1-R | CGTCGACACCGTGATTAGATTG |
| oEG141 | ChIP | TELVIIL-F | AGCCCGAGCCTGTACTAAAT |
| oEG142 | ChIP | TELVIIL-R | CAAAAGAAACTTTTCATGGCA |
